# Auditory beat perception is related to speech output fluency in post-stroke aphasia

**DOI:** 10.1101/2020.04.02.022475

**Authors:** James D. Stefaniak, Matthew A. Lambon Ralph, Blanca De Dios Perez, Timothy D. Griffiths, Manon Grube

## Abstract

Aphasia affects at least one third of stroke survivors, and there is increasing awareness that more fundamental deficits in auditory processing might contribute to impaired language performance in such individuals. We performed a comprehensive battery of psychoacoustic tasks assessing the perception of tone pairs and sequences across the domains of pitch, rhythm and timbre in 17 individuals with post-stroke aphasia and 17 controls. At the group level, we showed a significant difference in auditory perception in only one test (Dynamic Modulation detection). At the level of individual differences we demonstrated a correlation between metrical pattern (beat) perception and speech output fluency with strong effect (Spearman’s rho = 0.72). This was specific in terms of the auditory tests and dissociated from more basic auditory timing perception, which did not correlate with output fluency. This was also specific in terms of the language and cognitive measures, amongst which phonological, semantic and executive function did not correlate with beat detection. We interpret the data in terms of a requirement for the analysis of the metrical structure of sound to construct fluent output, with both being a function of higher-order “temporal scaffolding”. The beat perception task herein allows measurement of timing analysis without any need to account for motor output deficit, and could be a potential clinical tool to examine this. This work suggests strategies to improve fluency after stroke by training in metrical pattern perception.

## Introduction

Post-stroke aphasia (PSA) is prevalent (Engelter *et al.*, 2006) and debilitating (Tsouli *et al.*, 2009), and there is increasing awareness that deficits in non-verbal cognitive processing might contribute to language performance in such individuals (Robson *et al.*, 2013; Geranmayeh *et al.*, 2014; Schumacher *et al.*, 2019). Indeed, there is evidence from multiple aetiologies including congenital amusia (Foxton *et al.*, 2004; Foxton *et al.*, 2006), developmental dyslexia (Witton *et al.*, 1998), specific language impairment (Corriveau *et al.*, 2007), children who stutter (Wieland *et al.*, 2015) and primary progressive aphasia (Goll *et al.*, 2010; Grube *et al.*, 2016) that language impairment can co-occur with more fundamental deficits in auditory processing, just as auditory processing abilities associate with language performance in healthy children (Grube *et al.*, 2012) and adults (Grube *et al.*, 2013). However, there has been no systematic investigation of auditory processing deficits across the range of pitch, rhythm and timbre in post-stroke aphasia and whether these are associated with patients’ language impairments. In particular, it is unknown whether ‘output’ speech production deficits are associated with ‘input’ auditory processing deficits post stroke, as has been found in primary progressive aphasia (Goll *et al.*, 2010; Grube *et al.*, 2016). We have therefore assessed central auditory processing of pitch, rhythm and timbre in a cohort of individuals with chronic PSA.

There are several specific questions we sought to address.

The small number of previous studies assessing central auditory processing in PSA have failed to assess the full range of auditory processing domains pertinent to language, including pitch and melody, rhythm and timing, and timbre (Rosen, 1992; Chi *et al.*, 1999). For instance, both Fink *et al.* (2006) and Von Steinbüchel *et al.* (1999) found impaired temporal order judgement in PSA, but did not assess melody, rhythm or timbre processing (von Steinbüchel *et al.*, 1999; Fink *et al.*, 2006). Similarly, Robson *et al.* (2013, 2019) assessed pure tone frequency discrimination and timbre but not rhythm in a selective PSA group of Wernicke’s aphasia (Robson *et al.*, 2013; Robson *et al.*, 2019), while Robin *et al.* (1990) assessed pitch and rhythm but not timbre processing (Robin *et al.*, 1990). Furthermore, previous studies have tested the auditory processing of tone pairs or short tone sequences rather than longer and more abstract tone sequences, which may overlook processes used during the comprehension and production of more complex, naturalistic auditory stimuli such as language. We therefore sought to measure auditory processing of tone pairs and sequences across the range of pitch, rhythm and timbre domains in individuals with PSA.

Individuals with primary progressive aphasia have deficits in non-verbal auditory processing (Goll *et al.*, 2010). Intriguingly, recent work found that the non-fluent variant is particularly associated with auditory processing deficits for sequences of tones, suggesting the existence of a common neural substrate for processing auditory rhythmic structure in both auditory input and speech output (Grube *et al.*, 2016). Such a relationship has never been investigated in PSA; the aforementioned studies have only examined the relationship between auditory processing and language tests assessing comprehension but not speech production. Furthermore, previous work identified differences in auditory sequence processing at the group level between non-fluent and fluent variants of primary progressive aphasia (Grube *et al.*, 2016), but was not able to assess whether auditory sequence processing correlated with behavioural measures of speech fluency across individuals. If present, a relationship between auditory processing of tone sequences and speech output fluency would have significant implications for our understanding of fluent speech production, and for rehabilitation strategies such as melodic intonation therapy that might target such underlying auditory processing deficits in PSA (Zumbansen *et al.*, 2014). We therefore had the *a priori* hypothesis that speech output fluency would correlate with the auditory processing of sequences of tones, but not with the processing of simpler tone pairs, in PSA.

We performed a comprehensive battery of psychoacoustic tasks assessing pitch, rhythm and timbre in a cohort of individuals with chronic PSA for whom a large range of detailed language assessments encompassing phonology, semantics, fluency and executive function were available. Our aims were: a) to characterise the profile of auditory processing deficits in chronic PSA; and b) to test the hypothesis that speech output fluency would correlate with the ability to process sequences of tones, but not with the ability to process simpler tone pairs.

## Materials and methods

### Participants

17 individuals with chronic PSA (the ‘PSA subgroup’) and 17 healthy controls participated in the psychoacoustic testing. Individuals with PSA were at least one year post left hemispheric stroke (either ischaemic or haemorrhagic), pre-morbidly right-handed and part of a larger cohort of 76 stroke survivors recruited from community groups throughout the North West of England for whom extensive neuropsychological and imaging data was available. A number of the participants with PSA have been included in previous publications (Halai *et al.*, 2017; Zhao *et al.*, 2018). Controls were recruited from the volunteer panel of the MRC Cognition and Brain Sciences Unit, were right handed and had no history of neurological injury. All participants were native English speakers. Informed consent was obtained from all participants according to the Declaration of Helsinki under approval from the local research ethics committee.

Demographic and clinical variables for the PSA subgroup and controls are shown in Supplementary Table S1. Over half (52.9%) of the PSA subgroup had a classification of ‘anomia’; the remainder were classified as Broca’s aphasia (17.6%), conduction aphasia (17.6%), mixed nonfluent aphasia (5.9%) and transcortical motor aphasia (5.9%) (Supplementary Table S1). The PSA subgroup and controls were not statistically significantly different at the group level for age (mean 60.9 [SD 9.5] years in stroke survivors vs 62.5 (SD 4.3) in controls; *t*-test, t_22_=-0.60, two-sided p=0.55) and sex (8 females in stroke survivors vs 9 females in controls; Chi-Square test, χ^2^_1_=0.12, two-sided p=0.73) but the PSA subgroup had significantly fewer years of education than controls (median 11.0 [IQR 6.0] years in stroke survivors vs 16.0 [IQR 4.0] in controls; Mann-Whitney U-test, U=216.5, two-sided p=0.01).

Pure-tone audiograms were recorded in all participants using a Guymark Maico MA41 audiometer with Sennheiser HDA300 headphones to assess for evidence of peripheral hearing loss that might affect performance on the psychoacoustic tests. All participants had mean hearing levels < 20 dB HL between 0.25–1 kHz at octave intervals in at least one ear (Supplementary Table S2). There were no significant differences in pure tone audiometry thresholds between the PSA subgroup and controls (Supplementary Table S3). Mean pure tone detection thresholds between 0.25-1kHz were not significantly correlated with any of the psychoacoustic measures in the PSA subgroup (Supplementary Table S4).

### Lesion overlap map

Lesions were segmented from structural T_1_-weighted MRI images and normalised to MNI space using LINDA v0.5.0 (www.github.com/dorianps/LINDA) (Pustina *et al.*, 2016; Ito *et al.*, 2019). The lesion overlap map of the PSA subgroup is shown in Fig. 1; it encompasses most of the left hemisphere including subcortical white matter.

**Figure 1:**
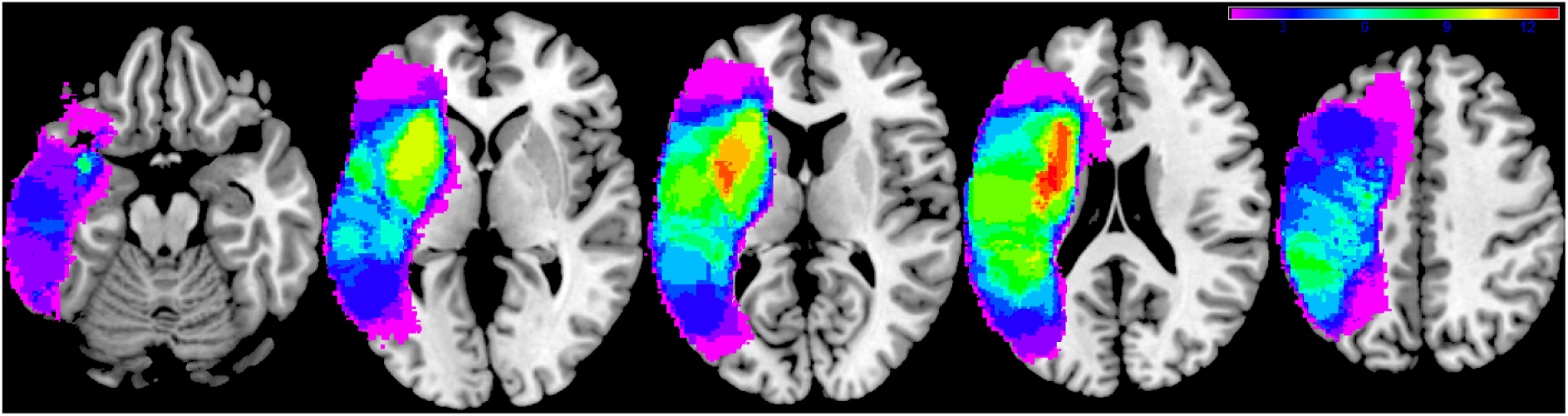
Lesion overlap map. Lesion overlap map for the 17 patient post-stroke aphasia subgroup in MNI space and in neurological convention. Colour bar represents overlap number between 1 and 13.

### Neuropsychological tests

The entire cohort of 76 stroke survivors had previously been administered an extensive battery of neuropsychological tests and principal component analysis (PCA) results from earlier collection rounds of this database have been published (Butler *et al.*, 2014; Halai *et al.*, 2017; Halai *et al.*, 2020 (in press)). Controls performed the ‘Cookie theft’ description (Boston Diagnostic Aphasia Examination (Goodglass and Kaplan, 1983)) and the Raven’s Coloured Progressive Matrices (Raven, 1962). Neuropsychological test scores for the entire cohort of 76 individuals with PSA are shown in Supplementary Table S5.

### Psychoacoustic tests of pitch, rhythm and timbre

The psychoacoustic tests used in this study had been developed and published previously (Grube *et al.*, 2016) and were administered using Matlab R2018a between February 2018 and July 2019. In both the PSA subgroup and controls we measured: pitch basic change detection (P1); pitch detection of local change (P2); pitch detection of global change (P3); single time interval discrimination (R1); isochrony deviation detection (R2); metrical pattern discrimination (R3); and Dynamic Modulation (DM) detection. P2 and P3 required a same-different choice at a fixed difficulty level, whereas the other tests (P1, R1, R2, R3, DM) used a two-alternative forced-choice adaptive paradigm with a two-down, one-up algorithm that estimated the 70.9% correct point of the psychometric function (Levitt, 1971). Of note, single time interval discrimination is taken to be a measure of rhythm processing in this study.

At the start of each test, instructions were explained in a way that was understandable for the participant, and practice trials with feedback were performed to ensure that the participant understood the task and could perform reliably at the easiest difficulty level. A basic description of the psychoacoustic tasks are provided below, but a more detailed description was published previously (Grube *et al.*, 2016).

All three pitch tasks (P1-P3) used pure tones of 250ms duration and a 2000ms interval between stimuli on each trial. In P1, two pairs of pure tones were presented on each trial. The participant had to indicate whether the first or the second pair had a change in frequency (either up or down). The size of the change in frequency was adaptively reduced and the threshold, in semitones, was used as the outcome measure. The task comprised of 50 trials.

In P2 and P3, two sequences of four tones were played on each trial. Tones within each sequence had varying pitch; participants had to indicate whether the first and second sequences were the same or different to each other. In P2, on the ‘different’ trials, there was one change in frequency in the third or fourth tone that produced a ‘local’ change in pitch without a change in the ‘global’ pattern of ‘ups and downs’. In P3, on the ‘different’ trials, the change in frequency of the third or fourth tone produced a change in the global pitch ‘contour’ of ‘ups and downs’. P2 and P3 each consisted of 40 trials (20 ‘same’ and 20 ‘different). The percentage (score) correct on each task was the outcome measure.

All three rhythm tasks (R1-R3) used 500Hz, 100ms pure tones. In R1, two tone pairs were played on each trial. Each pair of tones had a slightly different inter-onset interval; the range of inter-onset intervals was 300-600ms. Participants had to indicate whether the first or the second tone pair had the longer interval. R1 comprised of 50 trials; the difference between the interval of the first and the second pair was adjusted adaptively. The threshold, expressed as the percentage difference between the interval of the shorter and longer tone pair relative to the inter-onset-interval of the shorter pair, was used as the outcome measure.

In R2, each trial consisted of two five-tone sequences. One sequence was perfectly isochronous (i.e. had a constant inter-onset interval, with a value between 300 to 600ms that was varied between trial); the other sequence was not isochronous but had one lengthened inter-onset interval (between the third and fourth tone). Participants had to indicate whether the first or the second sequence contained an ‘extra gap’. R2 comprised of 50 trials; the difference between the lengthened inter-onset interval and the isochronous inter-onset interval was adaptively adjusted. The threshold, expressed as the percentage difference (relative to the otherwise isochronous inter-onset-interval), was used as the outcome measure.

In R3, each trial consisted of three seven-tone rhythmic sequences. Each sequence contained seven tones distributed over 16 time units of 180 to 220ms each, with the unit duration varied between sequences (i.e. within trials). In its correct version the pattern of the sequences featured a strongly metrical beat induced by accented tones occurring every fourth time unit (Grube and Griffiths, 2009). The first sequence on each trial always had the correct pattern, one of the second or the third sequence on each trial was the same as the first, but the remaining sequence (i.e. the second or the third) had a perturbation in the rhythm that affected the entire pattern and distorted its metricality (for details see (Grube and Griffiths, 2009)). The participant had to indicate whether the second or the third sequence sounded ‘different’ or ‘wrong’. R3 contained 50 trials; the percentage difference in relative interval timing was adaptively adjusted and the threshold, expressed as the percentage of perturbation relative to the correct pattern, used as the outcome measure as described in (Grube and Griffiths, 2009).

In the DM detection task, two 1000ms sounds were played on each trial. One sound was unmodulated, the other sound was spectro-temporally modulated (Chi *et al.*, 1999; Grube *et al.*, 2012). Participants had to indicate whether the first or the second sound was modulated. The degree of modulation was adaptively adjusted over 50 trials in the adaptive paradigm; the threshold was used as the outcome measure.

### Statistical analysis

All variables were assessed for normality using the Kolmogorov-Smirnov test. As the PSA subgroup had significantly fewer years of education than controls (see earlier), group level comparisons between PSA and control participants were performed with years of education as a covariate of no interest. As several psychoacoustic, neuropsychological and pure tone audiometry variables were not normally distributed, non-parametric tests (Mann-Whitney U tests, one-way rank analysis of covariance (ANCOVA) (Quade, 1967), Spearman correlations) were used.

Given the large number of neuropsychological measures available for our PSA subgroup and the whole cohort we recruited them from, we used a varimax-rotated PCA to reduce these scores to a smaller number of dimensions, as has been done previously (Butler *et al.*, 2014; Halai *et al.*, 2017). As PCA was unlikely to be stable in our subgroup of 17 PSA participants with psychoacoustic data (Preacher and MacCallum, 2002), we performed the PCA on the correlation matrix of neuropsychological test scores of the entire cohort of PSA participants (n=76), which has been shown formally to be highly reliable and stable (Halai *et al.*, 2020 (in press)). Scores from Principal Components (PCs) with an eigenvalue greater than 1 were taken to be estimates of underlying cognitive components in our 17 PSA participants and were used in correlation analyses with the psychoacoustic measures.

We defined statistical significance as p<0.05 with Bonferroni correction (by the number of tests) applied to the significance thresholds (i.e. reported p-values are uncorrected). Reported p-values for correlations between neuropsychological and/or psychoacoustic scores are one-tailed due to *a priori* hypotheses as to the direction of the associations, i.e. that better performance on auditory tasks would be associated with higher language scores.

### Data availability

The authors confirm that the data of this study are available upon reasonable request. See the online Supplementary Material for additional methodological details.

## Results

### Speech output fluency and executive function

At the group level, the PSA subgroup was not significantly worse than controls on the Raven’s Progressive Coloured Matrices (one-way rank ANCOVA with years of education as covariate, F(1,32)=0.41, p=0.53). The PSA subgroup produced significantly fewer words per minute (one-way rank ANCOVA with years of education as covariate, F(1,32)=23.35, p=0.00003), mean length of utterances (one-way rank ANCOVA with years of education as covariate, F(1,32)=24.75, p=0.00002) and speech tokens (one-way rank ANCOVA with years of education as covariate, F(1,32)=15.72, p=0.0004) on the ‘Cookie theft’ description task than controls (Table 1). This confirms that the PSA subgroup had significantly impaired fluency of connected speech, but relatively preserved executive function, compared to controls. Speech fluency and executive function test scores for the PSA subgroup and controls are shown in Supplementary Table S1.

**Table 1:** Group level comparisons of speech fluency and executive function between participants with post-stroke aphasia and controls. ‘P value’ corresponds to p-values from one-way rank ANCOVAs comparing neuropsychological scores between the post-stroke aphasia subgroup and controls, with years of education included as a covariate. All scores are expressed as a percentage of the maximum score in the sample of 17 participants with aphasia. * indicates the p-value is significant at the Bonferroni corrected significance threshold of p<0.0125 (corrected for 4 comparisons).

| Neuropsychological test | Participants with aphasia (median, IQR) | Controls (median, IQR) | P value |
| --- | --- | --- | --- |
| Raven’s Progressive Coloured Matrices | 88.89 (13.88) | 94.44 (16.67) | 0.53 |
| **Cookie Theft Description:** |  |  |  |
| Words Per Minute | 23.93 (31.39) | 68.37 (13.89) | 0.00003* |
| Mean Length of Utterance | 47.30 (33.76) | 127.55 (49.73) | 0.00002* |
| Number of speech tokens | 14.92 (11.00) | 50.16 (43.00) | 0.0004* |

### Auditory processing

Fig. 2 shows the time course of the patients’ performances on the seven psychoacoustic tests. P2 and P3 required a same-different choice at a fixed difficulty level and their cumulative scores correct graphs demonstrate a progressive increase of the cumulative correct score throughout both tests. The other psychoacoustic tests (P1, R1-R3, DM) used an adaptive difficulty paradigm and their “staircase plots” demonstrate a reliable decrease in the detection or discrimination threshold until a stable plateau was reached before the end of each test. Fig. 2 suggests that the psychoacoustic tests were performed reliably by patients in the PSA subgroup, without evidence of fatigue or losing track of the task. Psychoacoustic scores for the PSA subgroup and controls are shown in Supplementary Table S1.

**Figure 2:**
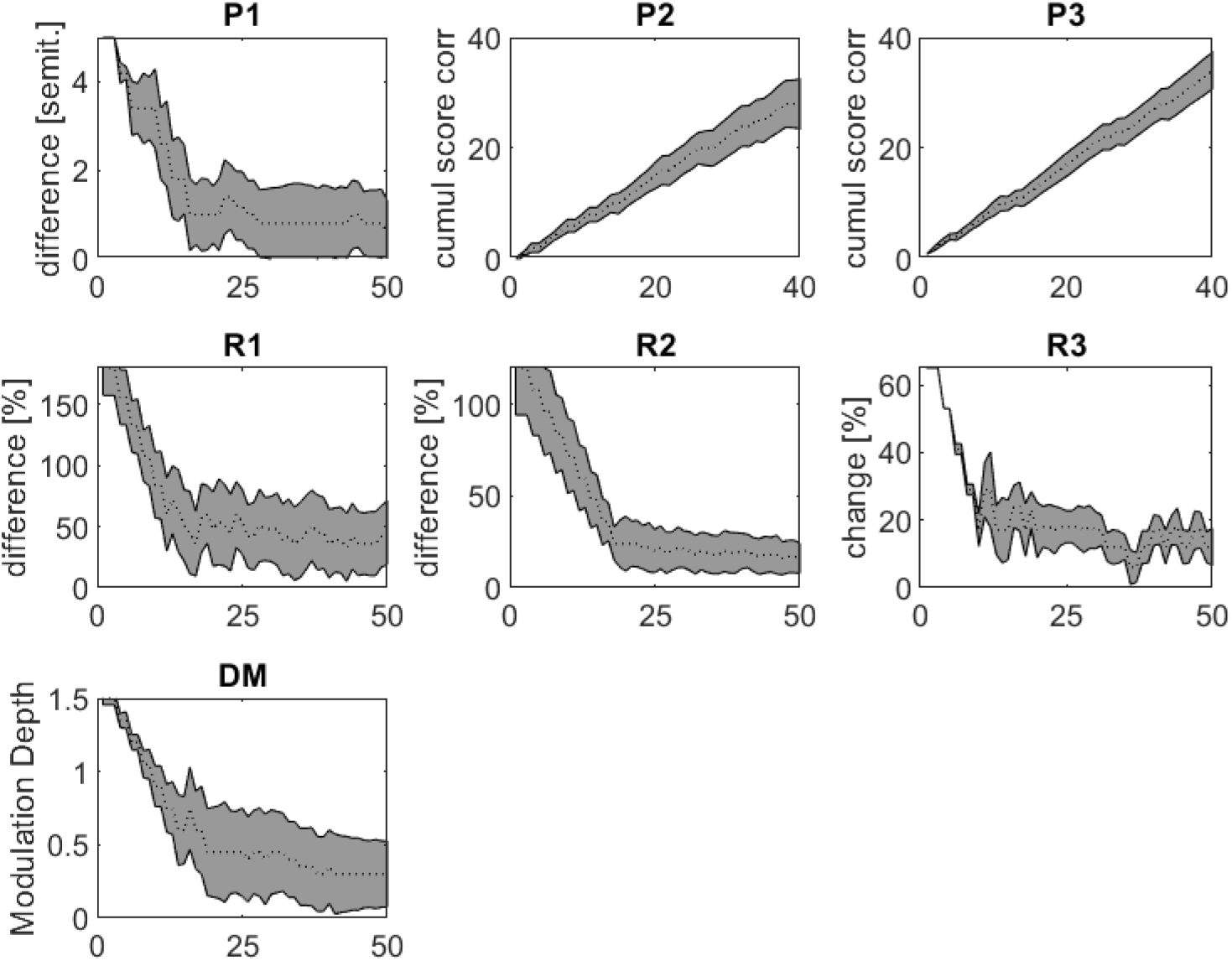
Reliable timecourses for the seven psychoacoustic tests. Each graph shows the timecourse of all patients’ performances throughout one of the seven psychoacoustic tests. X-axes represent trial number. Dotted lines are the median performances and the grey shaded areas represent ± the mean absolute difference from the median performance for all patients. Abbreviations: ‘P1’ = pitch basic change detection; ‘P2’ = pitch detection of local change; ‘P3’ = pitch detection of global change; ‘R1’ = rhythm single time interval discrimination; ‘R2’ = rhythm isochrony deviation detection; ‘R3’ = rhythm metrical pattern discrimination for a strongly metrical sequence; ‘DM’ = Dynamic Modulation detection.

The PSA group were only statistically significantly impaired in comparison to the controls on one of the seven psychoacoustic tests performed; this was Dynamic Modulation detection (one-way rank ANCOVA with years of education as covariate, F(1,32)=12.58, p=0.001; threshold higher in participants with PSA than controls) (Table 2). For the other six psychoacoustic tests performed there was no significant impairment at the group level (Table 2).

**Table 2:** Group level comparisons of psychoacoustic scores between participants with post-stroke aphasia and controls. ‘P value’ corresponds to p-values from one-way rank ANCOVAs comparing psychoacoustic scores between the post-stroke aphasia subgroup and control group, with years of education included as a covariate. For tests in which the outcome measure was a threshold, lower scores correspond to better performance. * indicates the p-value is significant at the Bonferroni corrected significance threshold of p<0.007 (corrected for 7 comparisons).

| Psychoacoustic measure | Participants with aphasia (median, IQR) | Controls (median, IQR) | P value |
| --- | --- | --- | --- |
| Pitch basic change detection threshold (semitones) | 0.83 (1.32) | 0.30 (0.13) | 0.03 |
| Pitch detection of local change (% correct) | 70.00 (19.00) | 85.00 (15.00) | 0.06 |
| Pitch detection of global change (% correct) | 85.00 (17.00) | 95.00 (7.00) | 0.06 |
| Rhythm single time interval discrimination threshold (%) | 44.50 (52.00) | 18.00 (24.00) | 0.12 |
| Rhythm isochrony deviation detection threshold (%) | 18.90 (15.50) | 13.00 (7.80) | 0.06 |
| Rhythm metrical pattern discrimination threshold (%) | 14.00 (6.00) | 11.00 (4.83) | 0.27 |
| Dynamic Modulation detection threshold (%) | 0.33 (0.48) | 0.15 (0.08) | 0.001* |

It was possible that PSA participants might have been significantly impaired on psychoacoustic tasks at the individual level, even if there was no group difference compared to controls. We therefore compared, individually, each PSA participant’s psychoacoustic scores to the control group data using the Bayesian Test for a Deficit controlling for years of education as a covariate (Crawford *et al.*, 2011). This conservative analysis identified a number of impairments at the individual level. In addition, several participants with PSA were unable to perform one or more of the rhythm processing tasks at the easiest difficulty level; we were not able to include these participants in group difference or correlation analyses of the corresponding psychoacoustic test as a reliable threshold was not able to be obtained, but these participants clearly had impaired auditory processing as well. Overall, five participants with PSA had significantly impaired performance on DM detection; three participants with PSA had significantly impaired performance on basic pitch discrimination (P1); one participant with PSA had significantly impaired performance on single time interval discrimination (R1) with an additional participant being unable to perform this task at the easiest difficulty level; two participants with PSA had significantly impaired performance on isochrony deviation detection (R2) with an additional three participants being unable to perform this task at the easiest difficulty level; and two participants with PSA were unable to perform metrical pattern discrimination (R3) at the easiest difficulty level (Supplementary Table S6). We did not find evidence that any of the participants with PSA were significantly impaired at the individual level on either of the psychoacoustic tests requiring processing of pitch in short tone sequences (P2, P3) (Supplementary Table S6).

### Principal Component Analysis of neuropsychological scores

We performed varimax-rotated PCA on the correlation matrix of neuropsychological test scores of the entire cohort of stroke survivors (n=76), including those who underwent psychoacoustic testing (n=17) and those who did not (n=59). The Kaiser-Meyer-Olkin value was 0.82, indicating adequate sampling (Kaiser, 1974). Four rotated PCs with eigenvalues greater than 1 explaining 27.73% (PC1), 17.71% (PC2), 17.63% (PC3) and 15.22% (PC4) of the variance were obtained, consistent with previous publications using data from this cohort of stroke survivors (Butler *et al.*, 2014; Halai *et al.*, 2017; Halai *et al.*, 2020 (in press)) (Table 3). PC1 was loaded onto primarily by immediate non-word and word repetition (subtests 8 and 9 from the Psycholinguistic Assessment of Language Processing in Aphasia battery (Kay *et al.*, 1992)), Boston Naming Test (Kaplan *et al.*, 1983), Forward Digit Span (Wechsler, 1987), and Cambridge Semantic Battery picture naming (Bozeat *et al.*, 2000); it was interpreted as representing phonological ability. PC2 was loaded onto primarily by the Cambridge Semantic Battery spoken word-to-picture matching (Bozeat *et al.*, 2000), spoken sentence comprehension from the Comprehensive Aphasia Test (Swinburn *et al.*, 2005), 96-trial synonym judgement test (Jefferies *et al.*, 2009) and Type-Token Ratio from the ‘Cookie theft’ picture description task; PC2 was interpreted as representing semantic processing. PC3 was loaded onto mainly by the number of speech tokens, Mean Length of Utterance and Words Per Minute from the ‘Cookie theft’ picture description task; this component was interpreted as representing fluency of connected speech. PC4 was loaded onto primarily by the Raven’s Coloured Progressive Matrices and the Brixton Spatial Anticipation Test (Burgess and Shallice, 1997); this was interpreting as representing executive function.

**Table 3:** Component matrix of neuropsychological scores from the entire cohort of participants with post-stroke aphasia. Varimax rotated principal component analysis was performed on the neuropsychological scores of the entire cohort of 76 individuals with post-stroke aphasia. The loading of each score onto each rotated principal component is shown. Variables with major loadings (defined as >0.50) are in bold. Abbreviation: ‘CSB’ = Cambridge Semantic Battery.

| Neuropsychological test | Component loadings |  |  |  |
| --- | --- | --- | --- | --- |
|  | PC 1 | PC 2 | PC 3 | PC 4 |
| Immediate non-word repetition | **0.90** | 0.09 | 0.19 | 0.19 |
| Immediate word repetition | **0.90** | 0.18 | 0.14 | 0.13 |
| Boston Naming Test | **0.83** | 0.38 | 0.16 | 0.11 |
| CSB Picture Naming | **0.84** | 0.40 | 0.12 | 0.13 |
| Forward Digit Span | **0.58** | 0.49 | 0.12 | 0.24 |
| CSB Spoken Word-to-Picture Matching | 0.24 | **0.77** | 0.15 | 0.21 |
| Spoken Sentence Comprehension | 0.39 | **0.53** | 0.20 | 0.46 |
| Synonym Judgement Test | 0.32 | **0.51** | 0.25 | 0.47 |
| ‘Cookie Theft’ Type-Token Ratio | 0.23 | **0.82** | -0.02 | -0.08 |
| ‘Cookie Theft’ number of speech tokens | -0.01 | -0.06 | **0.89** | 0.19 |
| ‘Cookie Theft’ Mean Length of Utterance | 0.25 | 0.21 | **0.88** | 0.12 |
| ‘Cookie Theft’ Words Per Minute | 0.25 | 0.14 | **0.79** | 0.12 |
| Raven’s Coloured Progressive Matrices | 0.12 | 0.11 | 0.10 | **0.86** |
| Brixton Spatial Anticipation Test | 0.16 | 0.05 | 0.20 | **0.83** |

### Principal Component 3 as a measure of speech output fluency

In the PCA performed on the neuropsychological scores of the entire cohort of 76 individuals with PSA, PC3 was loaded by measures of connected speech fluency obtained from the ‘Cookie theft’ description task (all loadings ≥0.79); PC3 was not loaded by any of the other neuropsychological scores (all other loadings ≤0.25) (Table 3). To ensure that PC3 remained associated with speech fluency but not the other neuropsychological scores in the PSA subgroup, we computed Spearman correlations between PC3 score and all 14 neuropsychological test scores in the PSA subgroup (Supplementary Table S7). The three ‘Cookie theft’ fluency scores (Words Per Minute, Mean Length of Utterance and number of speech tokens) correlated positively with PC3 (lowest Spearman rho 0.91), even after Bonferroni correcting for 14 multiple comparisons (Supplementary Table S7). By contrast, all other neuropsychological test scores did not significantly correlate with PC3 (highest Spearman rho 0.34), even before Bonferroni correction (Supplementary Table S7). This confirms that PC3 is a specific measure of speech output fluency in the PSA subgroup who underwent psychoacoustic testing.

In the PCA performed on the neuropsychological scores of the entire cohort of 76 individuals with PSA, PCs were, by definition, orthogonal. In the PSA subgroup, PC3 was neither significantly correlated with PC1 (Spearman’s rho=-0.25; two-sided p=0.33) nor with PC4 (Spearman’s rho=0.08; two-sided p=0.77) but was significantly negatively correlated with PC2 (Spearman’s rho=-0.67; two-sided uncorrected p=0.003; better PC3 fluency associated with worse PC2 semantics). In subsequent analyses assessing associations between auditory processing and PC3 (fluency), we therefore partialled out PC2 scores to ensure that any associations with PC3 could not be explained by confounding with PC2.

### Auditory sequence processing and speech output fluency

In order to test the hypothesis that the auditory analysis of tone sequences would be associated with speech output fluency in PSA, we computed Spearman correlations between psychoacoustic measures and PC3 (Table 4). The psychoacoustic measures that correlated significantly with PC3 after Bonferroni correction were: P3 (Spearman’s rho=0.63; uncorrected one-sided p=0.003; better pitch global change detection associated with better speech fluency); R1 (Spearman’s rho=-0.64; uncorrected one-sided p=0.004; better single time interval discrimination associated with better speech fluency); and R3 (Spearman’s rho=-0.72; uncorrected one-sided p=0.001; better metrical pattern discrimination associated with better speech fluency) (Fig. 3) (Table 4).

**Table 4:** Correlations between psychoacoustic measures and Principal Component 3 fluency. ‘P value’ corresponds to uncorrected one-sided p-values from Spearman correlations comparing psychoacoustic scores with PC3 fluency scores in the post-stroke aphasia subgroup. * indicates the p-value is significant at the Bonferroni corrected significance threshold of p<0.007 (corrected for 7 comparisons). Abbreviations: ‘PC’ = Principal Component.

| Psychoacoustic measure | Spearman correlation with PC3 |  |
| --- | --- | --- |
|  | Spearman’s rho | P value |
| Pitch basic change detection threshold (semitones) | -0.21 | 0.21 |
| Pitch detection of local change (% correct) | 0.43 | 0.04 |
| Pitch detection of global change (% correct) | 0.63 | 0.003* |
| Rhythm single time interval discrimination threshold (%) | -0.64 | 0.004* |
| Rhythm isochrony deviation detection threshold (%) | -0.20 | 0.24 |
| Rhythm metrical pattern discrimination threshold (%) | -0.72 | 0.001* |
| Dynamic Modulation detection threshold (%) | -0.15 | 0.29 |

**Figure 3:**
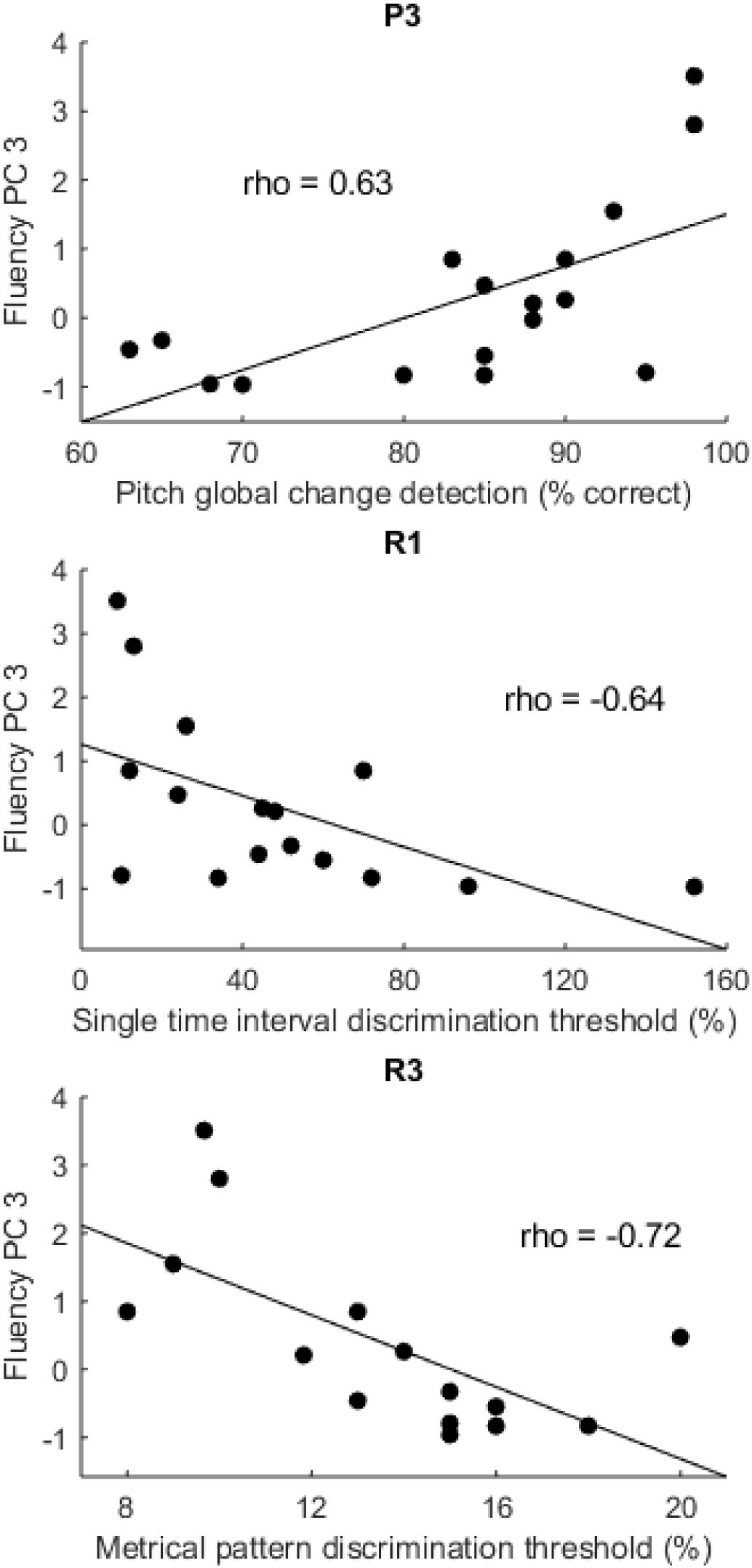
Psychoacoustic measures and speech fluency. Scatter plots showing significant correlations between ‘fluency’ Principal Component 3 scores (y-axis) and P3 (better pitch global change detection associated with better speech fluency), R1 (better single time interval discrimination associated with better speech fluency) and R3 (better metrical pattern discrimination associated with better speech fluency) (on x-axes) in the group of participants with post-stroke aphasia. The straight line represents the least-squares line of best fit. Abbreviations: ‘PC’ = Principal Component; ‘P3’ = pitch detection of global change; ‘R1’ = rhythm single time interval discrimination; ‘R3’ = rhythm metrical pattern discrimination for a strongly metrical sequence.

We wanted to identify psychoacoustic measures that were specifically associated with fluency, independent of other components of language. As PC3 scores were significantly negatively correlated with ‘semantic’ PC2 scores in this PSA subgroup, we repeated Spearman correlations between psychoacoustic measures and ‘fluency’ PC3 scores, while partialling out ‘semantic’ PC2 scores (Table 5). After partialling out PC2, only R3 remained significantly correlated with PC3 fluency scores (Spearman’s rho=-0.66, uncorrected one-sided p=0.005; more precise metrical rhythmic pattern discrimination associated with higher speech fluency) (Table 5). The psychoacoustic measures from the other tests based on tone sequences (P2, P3 and R2), or the more basic psychoacoustic tests using single sounds or tone pairs (P1, R1 and DM), were not significantly correlated with PC3 scores after partialling out PC2 and Bonferroni correcting for multiple comparisons (n=7; Table 5).

**Table 5:** Correlations between psychoacoustic measures and Principal Component 3 fluency, controlling for Principal Component 2. ‘P value’ corresponds to uncorrected one-sided p-values from Spearman correlations comparing psychoacoustic scores with PC3 fluency scores, partialling out PC2 scores, in the PSA subgroup. * indicates the p-value is significant at the Bonferroni corrected significance threshold of p<0.007 (corrected for 7 comparisons). Abbreviations: ‘PC’ = Principal Component.

| Psychoacoustic measure | Spearman correlation with PC3<br>(partialling out PC2) |  |
| --- | --- | --- |
|  | Spearman’s rho | P value |
| Pitch basic change detection threshold<br>(semitones) | -0.03 | 0.46 |
| Pitch detection of local change (% correct) | 0.23 | 0.20 |
| Pitch detection of global change (% correct) | 0.52 | 0.02 |
| Rhythm single time interval discrimination<br>threshold (%) | -0.27 | 0.17 |
| Rhythm isochrony deviation detection<br>threshold (%) | 0.27 | 0.81 |
| Rhythm metrical pattern discrimination<br>threshold (%) | -0.66 | 0.005* |
| Dynamic Modulation detection threshold (%) | 0.09 | 0.63 |

In order to confirm that rhythm metrical pattern discrimination (R3) was specifically associated with PC3 fluency, and not with other PCs of language, we performed Spearman correlations between R3 and PC1, PC2 and PC4 scores. R3 was not significantly correlated with PC1 (Spearman’s rho=-0.03; one-sided uncorrected p=0.45), PC2 (Spearman’s rho=0.39; one-sided uncorrected p=0.07) or PC4 (Spearman’s rho=0.29, one-sided uncorrected p=0.15). In order to confirm that R3’s association with PC3 was not due to increased executive processing demands during R3 relative to other psychoacoustic tasks, we performed Spearman correlations between R3 and PC3 while partialling out PC4 ‘executive’ scores. R3 remained significantly correlated with PC3 ‘fluency’ scores with no decrease in strength after partialling out PC4 (Spearman’s rho=-0.75; one-sided uncorrected p=0.001). Finally, to exclude effects of peripheral hearing, we confirmed that R3 was not significantly correlated with mean pure tone audiometry thresholds between 0.25-1kHz on either side (Supplementary Table S4).

### Dynamic Modulation detection and language

Since the PSA subgroup performed significantly worse than controls on DM detection (Table 2), we additionally performed Spearman correlations to look for any association between DM detection and the other PCs of language. DM detection was not significantly correlated with PC1 (Spearman’s rho=-0.03; one-sided uncorrected p=0.45), PC2 (Spearman’s rho=0.31; one-sided uncorrected p=0.11) or PC4 (Spearman’s rho=-0.14; one-sided uncorrected p=0.29). As DM detection has previously been associated with spoken comprehension in participants with Wernicke’s aphasia (Robson *et al.*, 2013), we additionally looked for associations between DM detection and individual neuropsychological tests assessing spoken comprehension. However, DM detection was not significantly correlated with Spoken Word-to-Picture Matching (Spearman’s rho=0.07; one-sided uncorrected p=0.39) or Spoken Sentence Comprehension (Spearman’s rho=0.09; one-sided uncorrected p=0.37).

### Summary

Group level comparisons identified significantly impaired fluent speech production and timbre DM processing in the PSA subgroup relative to controls. Individual case-group control comparisons identified further deficits in specific psychoacoustic measures. Correlations demonstrated a specific association between rhythm metrical pattern discrimination and ‘fluency’ Principal Component 3.

## Discussion

In order to investigate central auditory processing deficits in PSA, and whether auditory input processing of tone sequences relates to speech output fluency, we have performed an extensive battery of tests assessing the processing of pitch, rhythm and timbre in individuals with chronic aphasia following left hemisphere stroke and controls. Intriguingly, there was a strong and specific association between participants’ ability to discriminate the metrical pattern of strongly metrical tone sequences, and speech output fluency. This provides novel insights into the nature of fluent speech production in PSA and has conceptually replicated and extended previous work in primary progressive aphasia that suggested a relationship between auditory input processing of tone sequences and fluent speech production (Grube *et al.*, 2016).

A central aim of this work was to characterise central auditory processing deficits in PSA across the domains of pitch (P1-P3), rhythm (R1-R3) and timbre (DM detection). Previous work in PSA has tended to focus on auditory processing of single sounds or tone pairs (rather than sequences) in a limited number of domains, and has broadly found impaired processing of rhythm (Robin *et al.*, 1990; von Steinbüchel *et al.*, 1999; Fink *et al.*, 2006) and timbre (Robson *et al.*, 2013) following left hemisphere stroke despite relative preservation of pitch processing (Robin *et al.*, 1990; Robson *et al.*, 2013). The present study assessed pitch (P1-P3) and rhythm (R1-R3) of tone pairs and sequences, as well as an aspect of timbre processing (DM detection), in 17 individuals with a variety of aphasia profiles. Despite the PSA group having significantly impaired speech fluency (Table 1) and the lesion overlap map encompassing large parts of the left hemisphere (Fig. 1), we found no significant group-level impairment for the rhythm processing tasks performed (Table 2). We did not find evidence that any of the PSA participants were significantly impaired on the psychoacoustic tests processing pitch in tone sequences (P2, P3) (Supplementary Table S6). Three of the PSA participants had significantly impaired detection of basic pitch changes in tone pairs (P1); one participant had impaired discrimination of time intervals in tone pairs (R1) with an additional participant being unable to perform this task at the easiest difficulty level; two participants had impaired isochrony deviation detection in tone sequences (R2) with an additional three being unable to perform this task at the easiest difficult level; and two participants were unable to perform metrical pattern discrimination at the easiest difficult level (Supplementary Table S6). Timbre spectro-temporal modulation was the only psychoacoustic task that was significantly impaired in the PSA subgroup (Table 2), but we found no association between timbre processing ability and any of the PCs of language, despite an association with auditory comprehension having been demonstrated previously in Wernicke’s aphasia (Robson *et al.*, 2013; Robson *et al.*, 2019). A possible explanation for this relative lack of group-level auditory processing deficits is the heterogeneity of PSA and the possibility that auditory processing deficits differ depending on the lesion location and neuropsychological profile; indeed, our PSA subgroup did not contain anyone with classical Wernicke’s aphasia, as studied by Robson *et al.* (2013). We recruited individuals with any aphasia type, but the resultant subgroup consisted mainly of individuals with ‘expressive’ aphasia classifications (Supplementary Table S1). Previous studies associating timbre processing with auditory comprehension did so in a group selected for having Wernicke’s aphasia (Robson *et al.*, 2013; Robson *et al.*, 2019). It is possible that if our sample had included more individuals with severe comprehension deficits, we would have observed an association between timbre processing and auditory comprehension as well. Furthermore, three participants with PSA in this study were unable to perform one or more of the rhythm processing tasks at the easiest difficulty level (Supplementary Table S6). We were not able to include these participants in the group difference or correlation analyses because we could not quantify a reliable threshold; however, these three individuals with PSA clearly had impaired rhythm processing.

Our second main hypothesis was that there would be an association between the auditory processing of tone sequences and speech output fluency in PSA. This was based on previous research suggesting that individuals with the nonfluent variant of primary progressive aphasia are significantly impaired at auditory sequence processing relative to fluent variants (Grube *et al.*, 2016). The present study looked for associations between four tests of auditory sequence processing, as well as three psychoacoustic tests using tone pairs or sounds, and behavioural measures of speech output fluency. We found an association between speech output fluency (PC3) and one of the sequence processing tasks, namely rhythm metrical pattern discrimination (R3) (Table 5). Furthermore, this psychoacoustic measure was not correlated with the other PCs of language (PC1, PC2 or PC4) and thus was not a generic marker of aphasia severity. Neither the other sequence processing tasks (P2, P3, R2) nor the tasks involving processing of pairs of tones (P1, R1) or sounds (DM) were significantly associated with speech output fluency in this sample (Table 5). The present study therefore demonstrates that there is an association between ‘input’ auditory processing and ‘output’ speech fluency in PSA, but suggests that this association might specifically be between the discrimination of metrical pattern in tone sequences and fluent speech production. A novel implication of this study is therefore that the ability to detect metrical pattern in the incoming auditory stream might be a sensitive measure of an ability that is critical for the fluent production of connected speech.

The metricality of a tone sequence, as used in R3, is the higher-order temporal structure determined by the grouping of salvos of notes that induce the sense of a regularly occurring metrical ‘beat’ or ‘downbeat’, even when all notes have the same intensity, duration and pitch (Povel, 1984; Povel and Essens, 1985; Grube and Griffiths, 2009). A high degree of metrical structure enables us to anticipate and predict the higher-order temporal structure of upcoming sound, akin to a ‘temporal scaffolding’ based on the metrical beat (Grube and Griffiths, 2009). The association between metricality discrimination and fluency observed in this study suggests that the cognitive process of predicting the higher-order temporal structure of future sound might be important for both the discrimination of metricality in incoming auditory sound, and the production of metrical sound in fluent connected speech. Critically, isochrony deviation detection (R2) was not associated with speech fluency in this study. The difference between the two tasks is two-fold. Isochrony deviation detection (R2) tests lower-order differences in timing between consecutive tones in a simple isochronous sequence and uses a local deviation. By contrast, the deviation in the metrical task (R3) is distributed across the entire pattern, and the pattern is more abstract with a hierarchically organised beat structure. The lack of an association between isochrony deviation detection and fluency therefore suggests that the observed association between fluency and metrical pattern discrimination was not due to rhythm processing in general. Rather, it suggests an association between fluency and the ability to process the higher-order regularity of accented tones within a sequence as embodied by metrical patterns. This is in keeping with previous work showing an association between rapid automatised naming in healthy adults and their ability to detect a roughly regular beat in an otherwise irregular sequence, but not their ability to detect isochrony deviation (Bekius *et al.*, 2016).

It might be argued that the metrical pattern discrimination task (R3) is harder than the other three sequence processing tasks (P2, P3, and R2), because R3 requires comparisons between three sequences of tones (and the other three tasks require comparisons between two sequences of tones). However, we think this is unlikely to be the reason for the observed association between metrical pattern discrimination and speech fluency. Firstly, R3 requires the detection of a perturbation that is detectable within the sequence, unlike P2 or P3. Although R2 also requires the detection of a perturbation within the sequence, in R3 the deviation is distributed across the entire sequence with a number of different interval ratios, whereas R2 requires the analysis of one deviating interval from an otherwise isochronous beat. Secondly, the same sequence is used as the reference on all 50 trials in R3; a different metrical pattern does not have to be remembered on each trial. This is similar to R2 (which uses the same sequence at different tempi) but is in stark contrast to the two pitch sequence tasks (P2 and P3), which use different reference sequences on each trial that have to be remembered and compared to the target sequence. Thirdly, unlike P2 and P3, R3 used a two-alternative forced-choice adaptive difficulty paradigm with a two-down, one-up algorithm that is designed to reduce working memory load (Levitt, 1971). Fourthly, R3 was not significantly correlated with PC4 ‘executive’ scores, and its association with PC3 ‘fluency’ scores remained after partialling out PC4 ‘executive’ scores.

The strong and extremely robust correlation between rhythm metrical pattern discrimination and speech output fluency might seem surprising given the absence of impairments at the group and individual-level for this psychoacoustic task in the PSA subgroup relative to controls. Assuming that the impaired speech fluency observed in the PSA subgroup was a consequence of stroke, this suggests that some of the observed correlation between fluency and R3 might not be a consequence of stroke damaging a single neural substrate that is responsible for both fluency and R3. One possible explanation is that premorbid inter-individual differences in the ability to discriminate metrical pattern mitigates the effect of stroke on speech output fluency. Alternatively, recovery mechanisms post-stroke might have involved improvements in metricality discrimination which in turn helped fluency to recover. Similar possibilities were recently proposed when structural changes outside the lesion mask (and thus not directly caused by stroke) were associated with reading recovery in post-stroke central alexia (Aguilar *et al.*, 2018). It would be of great interest if rehabilitation strategies that use singing to aid recovery of propositional speech, such as melodic intonation therapy (Zumbansen *et al.*, 2014), were found to be mediated by improved metricality discrimination. Existing techniques that augment metricality using rhythmic auditory stimuli improve speech output post stroke (Brendel and Ziegler, 2008; Stahl *et al.*, 2011). These findings go further by suggesting that a potential avenue for future therapies might be to target auditory metricality discrimination to aid fluency in patients.

A limitation of the current work is that we were not able to elucidate the neural structures associated with metricality discrimination in PSA. Parkinson’s disease, in which basal ganglia degeneration occurs, is associated with impaired metricality-based rhythm discrimination (Grahn and Brett, 2009). Metrical rhythms elicit greater activation in the basal ganglia and supplementary motor area (Grahn and Brett, 2007), and striatal activation is thought to represent metricality prediction (rather than detection) (Grahn and Rowe, 2013). Future work should elucidate whether the striatum and supplementary motor area contribute to speech fluency through metricality discrimination in PSA.

## Conclusion

Our findings demonstrate that there is a strong and specific association between the ability to analyse metrical pattern structure in the incoming auditory stream and fluent production of connected speech. Critically, this association was not due to the ability to detect lower-order differences in timing between consecutive tones in a sequence. This has significant implications for our understanding of fluent speech production, and for rehabilitation strategies that might use rhythm processing to aid recovery of fluency post stroke.

## Acknowledgements

We would like to thank the volunteers for participating in this research.

