## Supplementary File for "Auditory beat perception is related to speech output fluency in post-stroke aphasia"

Supplementary material

Supplementary Table S1: Demographic, neuropsychological and psychoacoustic variables in participants with post-stroke aphasia and controls

Participants are ordered alphabetically within each group. Abbreviations: NA = Not Applicable. Neuropsychological test abbreviations: ‘Raven’s’ = Raven’s Coloured Progressive Matrices; ‘Words Per Minute’ = Words Per Minute on the BDAE Cookie Theft description; ‘Mean Length of Utterance’ = Mean Length of Utterance on the BDAE Cookie Theft description; ‘Speech Tokens’ = number of speech tokens on the BDAE Cookie Theft description’. Psychoacoustic test abbreviations: P1 = pitch basic change detection threshold (semitones); P2 = pitch detection of local change (% correct); P3 = pitch detection of global change (% correct); R1 = rhythm single time interval discrimination threshold (%); R2 = rhythm isochrony deviation detection threshold (%); R3 = rhythm metrical pattern discrimination threshold for a strongly metrical sequence (%); DM = Dynamic Modulation detection threshold (%). All neuropsychological scores were expressed as a percentage of the maximum score in the sample of 17 participants with aphasia who underwent psychoacoustic testing. All psychoacoustic scores were expressed as raw scores.

| **Identifier** |  | | |  | **Neuropsychological measures** | | | | **Psychoacoustic measures** | | | | | | |
| --- | --- | --- | --- | --- | --- | --- | --- | --- | --- | --- | --- | --- | --- | --- | --- |
|  | **Age** | **Sex** | **Years education** | **Classification** | **Raven’s** | **Words Per Minute** | **Mean Length of Utterance** | **Speech Tokens** | **P1** | **P2** | **P3** | **R1** | **R2** | **R3** | **DM** |
| **Healthy controls** |  |  |  |  |  |  |  |  |  |  |  |  |  |  |  |
| AB | 65 | F | 16 | NA | 80.6 | 52.6 | 57.1 | 31.1 | 0.33 | 100 | 98 | 39.0 | 32.0 | 18.0 | 0.13 |
| ACJ | 62 | M | 17 | NA | 94.4 | 75.1 | 127.6 | 32.7 | 0.27 | 98 | 98 | 11.0 | 0.7 | 13.0 | 0.14 |
| AT | 65 | M | 20 | NA | 100.0 | 61.5 | 104.6 | 33.7 | 0.20 | 93 | 98 | 14.0 | 6.3 | 7.3 | 0.22 |
| AW | 67 | M | 13 | NA | 94.4 | 77.5 | 201.5 | 63.2 | 0.40 | 85 | 90 | 14.0 | 10.0 | 7.0 | 0.20 |
| BM | 63 | F | 16 | NA | 100.0 | 92.1 | 131.8 | 86.3 | 0.30 | 80 | 98 | 15.0 | 14.3 | 15.0 | 0.17 |
| CJ | 57 | F | 16 | NA | 97.2 | 63.3 | 104.9 | 50.2 | 0.27 | 95 | 98 | 9.0 | 7.0 | 10.0 | 0.08 |
| JC | 62 | F | 16 | NA | 100.0 | 70.5 | 141.4 | 107.0 | 0.53 | 85 | 95 | 41.0 | 13.0 | 9.0 | 0.13 |
| JD | 57 | F | 17 | NA | 91.7 | 65.4 | 109.7 | 46.0 | 0.30 | 85 | 93 | 8.4 | 13.7 | 6.0 | 0.15 |
| JN | 56 | M | 12 | NA | 100.0 | 67.2 | 137.8 | 66.0 | 0.27 | 85 | 93 | 4.0 | 6.3 | 8.3 | 0.15 |
| LB | 66 | F | 11 | NA | 80.6 | 98.4 | 165.3 | 42.9 | 0.63 | NA | NA | 4.2 | 6.4 | 13.0 | 0.12 |
| NG | 61 | M | 20 | NA | 100.0 | 79.8 | 216.1 | 203.2 | 0.27 | 60 | 95 | 18.0 | 13.0 | 11.0 | 0.08 |
| NM | 56 | M | 13 | NA | 69.4 | 56.9 | 107.1 | 47.0 | NA | 80 | 88 | 29.0 | 14.7 | 14.0 | 0.43 |
| RR | 67 | M | 19 | NA | 97.2 | 65.5 | 143.6 | 154.3 | 2.65 | 65 | 53 | 57.6 | 13.0 | 19.0 | 0.21 |
| SB | 69 | F | 16 | NA | 77.8 | 78.9 | 250.0 | 66.0 | 0.40 | 100 | 98 | 22.5 | 15.7 | 11.0 | 0.10 |
| SM | 64 | F | 13 | NA | 86.1 | 68.4 | 89.3 | 35.9 | 0.33 | 73 | 90 | 52.0 | 15.7 | 11.0 | 0.19 |
| SP | 58 | M | 16 | NA | 97.2 | 65.3 | 67.1 | 46.7 | 0.27 | 88 | 95 | 19.5 | 13.3 | 11.0 | 0.12 |
| SW | 67 | F | 13 | NA | 94.4 | 73.0 | 120.3 | 79.0 | 0.20 | 95 | 100 | 18.0 | 10.0 | 11.0 | 0.17 |
| **Participants with aphasia** |  |  |  |  |  |  |  |  |  |  |  |  |  |  |  |
| AG | 62 | M | 11 | Broca | 75.0 | 8.5 | 37.7 | 9.5 | 0.93 | 65 | 85 | 34.0 | 18.8 | 16.0 | 0.28 |
| AL | 55 | F | 12 | Anomia | 91.7 | 100.0 | 60.3 | 19.0 | 0.60 | 70 | 93 | 26.0 | 32.0 | 9.0 | 0.34 |
| BH | 72 | M | 11 | Mixed nonfluent | 66.7 | 23.9 | 41.6 | 12.1 | 0.83 | 68 | 65 | 52.0 | 19.3 | 15.0 | 0.66 |
| CH | 46 | F | 13 | Anomia | 91.7 | 11.2 | 34.8 | 12.1 | 0.87 | 100 | 95 | 10.0 | 10.0 | 15.0 | 0.25 |
| DF | 51 | F | 11 | Anomia | 88.9 | 23.4 | 50.9 | 14.9 | 3.80 | 65 | 88 | NA | NA | NA | 0.53 |
| DM | 56 | M | 17 | Broca | 91.7 | 15.4 | 34.9 | 12.1 | 0.17 | 73 | 85 | 60.0 | 13.3 | 16.0 | 0.14 |
| Ebo | 47 | M | 11 | Anomia | 97.2 | 26.4 | 75.4 | 17.8 | 0.27 | 95 | 90 | 45.0 | 16.0 | 14.0 | 0.27 |
| GP | 63 | M | 11 | Anomia | 97.2 | 26.6 | 60.6 | 29.8 | 0.80 | 80 | 85 | 24.0 | 16.0 | 20.0 | 1.31 |
| JS | 73 | F | 19 | Anomia | 100.0 | 50.1 | 100.0 | 100.0 | 0.60 | 80 | 98 | 9.0 | 19.0 | 9.7 | 0.18 |
| MAD | 60 | F | 11 | Anomia | 83.3 | 17.6 | 23.8 | 7.3 | 0.60 | 57 | 68 | 96.0 | 63.3 | 15.0 | 0.80 |
| MH | 69 | M | 11 | Conduction | 80.6 | 51.9 | 47.3 | 17.5 | 2.43 | 83 | 90 | 70.0 | 22.7 | 13.0 | 1.11 |
| NC | 55 | M | 17 | Anomia | 88.9 | 42.4 | 80.5 | 22.2 | 0.90 | 53 | 83 | 12.0 | 12.0 | 8.0 | 0.15 |
| PBL | 49 | F | 16 | Conduction | 97.2 | 13.0 | 16.1 | 12.1 | 0.57 | 68 | 80 | 72.0 | NA | 18.0 | 0.25 |
| PR | 76 | F | 11 | Transcortical motor | 80.6 | 9.3 | 24.0 | 7.9 | 4.40 | 60 | 70 | 152.0 | NA | NA | 1.18 |
| RH | 68 | M | 17 | Conduction | 83.3 | 44.6 | 96.8 | 64.4 | 0.33 | 95 | 98 | 13.0 | 6.3 | 10.0 | 0.33 |
| ST | 67 | F | 11 | Broca | 50.0 | 9.4 | 33.6 | 9.5 | 1.77 | 60 | 63 | 44.0 | 43.3 | 13.0 | 0.53 |
| WE | 67 | M | 10 | Anomia | 91.7 | 26.1 | 58.9 | 21.9 | 2.03 | 80 | 88 | 48.0 | 27.3 | 11.8 | 0.33 |

Supplementary Table S2: Pure tone audiograms of participants with post-stroke aphasia and controls

Participants are ordered alphabetically within each group. Abbreviations: ‘dB’ = decibels; ‘Hz’ = Hertz.

|  | **Left ear hearing level (dB)** | | | | | **Right ear hearing level (dB)** | | | | |
| --- | --- | --- | --- | --- | --- | --- | --- | --- | --- | --- |
| **Identifier** | **250Hz** | **500Hz** | **1000Hz** | **2000Hz** | **4000Hz** | **250Hz** | **500Hz** | **1000Hz** | **2000Hz** | **4000Hz** |
| **Healthy controls** |  |  |  |  |  |  |  |  |  |  |
| AB | 0 | 5 | 15 | 15 | 30 | 5 | 5 | 10 | 15 | 25 |
| ACJ | 0 | 10 | 10 | 10 | 15 | 0 | 5 | 10 | 5 | 15 |
| AT | 10 | 15 | 10 | 30 | 40 | 10 | 10 | 15 | 10 | 35 |
| AW | 15 | 30 | 10 | 15 | 35 | 20 | 20 | 15 | 10 | 35 |
| BM | 15 | 15 | 10 | 15 | 30 | 15 | 10 | 15 | 20 | 25 |
| CJ | 10 | 10 | 5 | 10 | 20 | 20 | 20 | 15 | 15 | 30 |
| JC | 15 | 20 | 10 | 20 | 35 | 15 | 15 | 15 | 10 | 15 |
| JD | 10 | 15 | 10 | 15 | 30 | 15 | 15 | 20 | 25 | 25 |
| JN | 0 | 5 | 10 | 5 | 25 | 0 | 5 | 5 | 5 | 15 |
| LB | 15 | 15 | 15 | 20 | 30 | 15 | 15 | 20 | 5 | 35 |
| NG | 15 | 15 | 15 | 15 | 30 | 35 | 15 | 10 | 15 | 30 |
| NM | 25 | 20 | 20 | 30 | 40 | 15 | 15 | 10 | 15 | 30 |
| RR | 15 | 20 | 10 | 35 | 65 | 10 | 10 | 15 | 20 | 55 |
| SB | 5 | 15 | 15 | 10 | 25 | 5 | 10 | 15 | 15 | 25 |
| SM | 10 | 15 | 10 | 25 | 45 | 10 | 15 | 10 | 30 | 40 |
| SP | 10 | 10 | 10 | 15 | 35 | 5 | 10 | 10 | 10 | 30 |
| SW | 15 | 15 | 10 | 15 | 20 | 5 | 15 | 15 | 10 | 20 |
| **Participants with aphasia** |  |  |  |  |  |  |  |  |  |  |
| AG | -10 | -5 | -5 | 15 | 30 | -5 | 5 | 10 | 20 | 45 |
| AL | 20 | 25 | 35 | 30 | 15 | 15 | 15 | 30 | 30 | 20 |
| BH | 15 | 20 | 40 | 55 | 75 | 10 | 20 | 20 | 20 | 45 |
| CH | 15 | 20 | 15 | 10 | 10 | 0 | 0 | 10 | 5 | 20 |
| DF | 30 | 30 | 30 | 50 | 65 | 15 | 15 | 10 | 25 | 25 |
| DM | 0 | 5 | 0 | 5 | 20 | 5 | 5 | 15 | 5 | 25 |
| Ebo | 10 | 10 | 5 | 10 | 10 | 20 | 10 | 5 | 5 | 15 |
| GP | -5 | -5 | 0 | 25 | 55 | 0 | 0 | 5 | 25 | 55 |
| JS | 10 | 10 | 25 | 20 | 35 | 5 | 10 | 10 | 15 | 25 |
| MAD | 0 | 10 | 20 | 40 | 60 | -5 | 5 | 20 | 45 | 50 |
| MH | -5 | -5 | 5 | 10 | 40 | 0 | 5 | 5 | 10 | 25 |
| NC | 0 | 10 | 10 | 10 | 55 | 0 | 0 | 0 | 25 | 55 |
| PBL | 0 | 10 | 10 | 5 | 15 | 5 | 10 | 10 | 5 | 5 |
| PR | 25 | 30 | 25 | 25 | 50 | 20 | 25 | 15 | 25 | 55 |
| RH | 5 | 10 | 0 | 5 | 25 | 5 | 5 | 5 | 10 | 35 |
| ST | -5 | 0 | 10 | 20 | 50 | 10 | 15 | 35 | 40 | 55 |
| WE | 0 | -5 | 10 | 55 | 60 | 10 | 5 | 5 | 25 | 55 |

Supplementary Table S3: Group level comparisons of pure tone audiogram thresholds between participants with post-stroke aphasia and controls

‘P value’ corresponds to uncorrected two-sided p-values from Mann-Whitney U-tests comparing pure tone audiometry thresholds between participants with post-stroke aphasia and healthy controls. None of the p-values are significant at the Bonferroni corrected significance threshold of p<0.004 (corrected for 12 comparisons). Abbreviations: ‘dB’ = decibels; ‘Hz’ = Hertz; ‘IQR’ = interquartile range.

| **Pure Tone Freqency (Hz)** | **Threshold in participants with aphasia (median, IQR) (dB)** | **Threshold in healthy controls (median, IQR) (dB)** | **P value** |
| --- | --- | --- | --- |
| **Left ear** |  |  |  |
| 250 | 0.0 (18.0) | 10.0 (8.0) | 0.11 |
| 500 | 10.0 (23.0) | 15.0 (8.0) | 0.13 |
| 1000 | 10.0 (23.0) | 10.0 (5.0) | 0.95 |
| 2000 | 20.0 (25.0) | 15.0 (10.0) | 0.79 |
| 4000 | 40.0 (40.0) | 30.0 (13.0) | 0.39 |
| Mean 250-1000 | 6.7 (19.2) | 11.7 (5.8) | 0.22 |
| **Right ear** |  |  |  |
| 250 | 5.0 (13.0) | 10.0 (10.0) | 0.09 |
| 500 | 5.0 (10.0) | 15.0 (5.0) | 0.08 |
| 1000 | 10.0 (13.0) | 15.0 (5.0) | 0.25 |
| 2000 | 20.0 (18.0) | 15.0 (8.0) | 0.18 |
| 4000 | 35.0 (33.0) | 30.0 (13.0) | 0.27 |
| Mean 250-1000 | 8.3 (11.7) | 11.7 (7.5) | 0.11 |

Supplementary Table S4: Correlations between pure tone audiogram thresholds and psychoacoustic scores

Matrix showing Spearman correlations between mean pure tone audiometry thresholds between 250-1000Hz (columns) and psychoacoustic scores (rows) for the 17 participants with post-stroke aphasia who performed psychoacoustic tests. Each cell contains the rho and uncorrected, two-sided p-value in the form: rho (p value). None of the p-values are significant at the Bonferroni corrected significance threshold of p<0.007 (corrected for 7 comparisons). Abbreviation: ‘Hz’ = Hertz.

| **Psychoacoustic test** | **Left mean threshold 250-1000Hz** | **Right mean threshold 250-1000Hz** |
| --- | --- | --- |
| Pitch basic change detection threshold | 0.07 (0.80) | 0.03 (0.93) |
| Pitch detection of local change | -0.16 (0.55) | -0.24 (0.37) |
| Pitch detection of global change | 0.07 (0.80) | -0.25 (0.34) |
| Rhythm single time interval discrimination threshold | 0.03 (0.91) | 0.38 (0.15) |
| Rhythm isochrony deviation detection threshold | 0.08 (0.79) | 0.51 (0.06) |
| Rhythm metrical pattern discrimination threshold | -0.35 (0.20) | -0.12 (0.68) |
| Dynamic Modulation detection threshold | 0.03 (0.91) | 0.18 (0.48) |

Supplementary Table S5: Neuropsychological test scores for the entire cohort of 76 individuals with post stroke aphasia

Participants are ordered alphabetically. Abbreviations: ‘ID’ = Participant Identifier; ‘Nonword Repetition’ = immediate non-word repetition from the Psycholinguistic Assessment of Language Processing in Aphasia subtest 8; ‘Word Repetition’ = immediate word repetition from the Psycholinguistic Assessment of Language Processing in Aphasia subtest 9; ‘BNT’ = Boston Naming Test; ‘CSB Word-Picture Matching’ = Cambridge Semantic Battery spoken word-to-picture matching; ‘CSB Naming’ = Cambridge Semantic Battery picture naming; ‘Spoken Sentence Comprehension’ = spoken sentence comprehension subtest from the Comprehensive Aphasia Test; ‘Synonym Judgement’ = 96-trial synonym judgement test; ‘Raven’s’ = Raven’s Coloured Progressive Matrices; ‘Brixton’ = Brixton Spatial Anticipation Test; ‘Words Per Minute’ = Words Per Minute on the BDAE Cookie Theft description; ‘Type-Token Ratio’ = Type-Token Ratio on the BDAE Cookie Theft description; ‘Mean Length of Utterance’ = Mean Length of Utterance on the BDAE Cookie Theft description; ‘Speech Tokens’ = number of speech tokens on the Boston Diagnostic Aphasia Examination (BDAE) Cookie Theft description’. All scores are expressed as a percentage of the maximum score in this sample of 76 patients.

| **ID** | **Nonword Repetition** | **Word Repetition** | **BNT** | **Forward Digit Span** | **CSB Word-Picture Matching** | **CSB Naming** | **Spoken Sentence Comprehension** | **Synonym Judgement** | **Raven’s** | **Brixton** | **BDAE ‘Cookie Theft’ description** | | | |
| --- | --- | --- | --- | --- | --- | --- | --- | --- | --- | --- | --- | --- | --- | --- |
|  |  |  |  |  |  |  |  |  |  |  | **Words Per Minute** | **Type-Token Ratio** | **Mean Length of Utterance** | **Speech Tokens** |
| AB | 26.7 | 77.5 | 28.3 | 37.5 | 95.3 | 53.1 | 75.0 | 75.0 | 88.9 | 88.9 | 26.6 | 51.6 | 84.3 | 38.7 |
| AD | 23.3 | 57.5 | 18.3 | 75.0 | 96.9 | 37.5 | 68.8 | 83.3 | 63.9 | 30.9 | 12.0 | 60.0 | 20.4 | 7.9 |
| Adr | 0.0 | 31.3 | 3.3 | 0.0 | 87.5 | 10.9 | 78.1 | 0.0 | 47.2 | 41.8 | 18.2 | 53.2 | 41.8 | 24.4 |
| AG | 73.3 | 77.5 | 78.3 | 100.0 | 100.0 | 85.9 | 87.5 | 89.6 | 75.0 | 56.4 | 8.5 | 70.0 | 37.7 | 9.5 |
| AL | 90.0 | 100.0 | 85.0 | 87.5 | 100.0 | 92.2 | 84.4 | 93.8 | 91.7 | 60.0 | 100.0 | 75.0 | 60.3 | 19.0 |
| AS | 0.0 | 35.0 | 0.0 | 25.0 | 76.6 | 6.3 | 50.0 | 42.7 | 47.2 | 47.3 | 8.5 | 33.3 | 10.2 | 1.9 |
| BH | 86.7 | 100.0 | 63.3 | 62.5 | 98.4 | 93.8 | 78.1 | 83.3 | 66.7 | 67.3 | 23.9 | 68.4 | 41.6 | 12.1 |
| BH | 23.3 | 48.8 | 15.0 | 62.5 | 100.0 | 39.1 | 87.5 | 90.6 | 83.3 | 65.5 | 7.9 | 80.0 | 14.8 | 7.9 |
| BS | 3.3 | 5.0 | 1.7 | 0.0 | 92.2 | 4.7 | 31.3 | 77.1 | 91.7 | 38.2 | 34.2 | 55.2 | 33.1 | 9.2 |
| CF | 33.3 | 70.0 | 11.7 | 25.0 | 89.1 | 29.7 | 68.8 | 77.1 | 91.7 | 50.9 | 3.8 | 100.0 | 8.9 | 3.5 |
| CH | 60.0 | 92.5 | 56.7 | 50.0 | 100.0 | 79.7 | 84.4 | 87.5 | 91.7 | 76.4 | 11.2 | 60.5 | 34.8 | 12.1 |
| DB | 70.0 | 85.0 | 8.3 | 37.5 | 64.1 | 7.8 | 31.3 | 59.4 | 86.1 | 40.0 | 11.3 | 33.3 | 44.9 | 38.1 |
| DBb | 0.0 | 36.3 | 0.0 | 25.0 | 59.4 | 0.0 | 12.5 | 49.0 | 30.6 | 38.2 | 41.2 | 46.9 | 41.8 | 10.2 |
| DC | 0.0 | 0.0 | 0.0 | 37.5 | 100.0 | 1.6 | 75.0 | 78.1 | 88.9 | 76.4 | 49.8 | 48.0 | 96.8 | 46.3 |
| DCS | 40.0 | 70.0 | 43.3 | 62.5 | 100.0 | 67.2 | 93.8 | 91.7 | 100.0 | 81.8 | 9.7 | 80.6 | 48.4 | 9.8 |
| DF | 53.3 | 93.8 | 46.7 | 37.5 | 100.0 | 76.6 | 62.5 | 78.1 | 88.9 | 43.6 | 23.4 | 72.3 | 50.9 | 14.9 |
| DL | 50.0 | 86.3 | 81.7 | 62.5 | 100.0 | 93.8 | 100.0 | 95.8 | 94.4 | 74.5 | 23.4 | 51.1 | 69.4 | 29.8 |
| DM | 60.0 | 70.0 | 70.0 | 37.5 | 98.4 | 71.9 | 56.3 | 95.8 | 91.7 | 50.9 | 15.4 | 73.7 | 34.9 | 12.1 |
| DR | 53.3 | 90.0 | 3.3 | 37.5 | 62.5 | 14.1 | 46.9 | 46.9 | 83.3 | 65.5 | 15.5 | 29.3 | 26.3 | 18.4 |
| DS | 56.7 | 88.8 | 75.0 | 50.0 | 100.0 | 81.3 | 87.5 | 93.8 | 72.2 | 72.7 | 22.5 | 87.0 | 68.8 | 7.3 |
| EB | 83.3 | 100.0 | 56.7 | 62.5 | 98.4 | 76.6 | 71.9 | 94.8 | 100.0 | 80.0 | 51.3 | 58.4 | 67.8 | 39.7 |
| EB | 66.7 | 81.3 | 31.7 | 50.0 | 98.4 | 70.3 | 75.0 | 69.8 | 66.7 | 47.3 | 26.9 | 68.0 | 75.4 | 17.8 |
| Ebo | 100.0 | 100.0 | 53.3 | 50.0 | 100.0 | 85.9 | 87.5 | 90.6 | 97.2 | 69.1 | 26.4 | 66.1 | 75.4 | 17.8 |
| ER | 53.3 | 70.0 | 63.3 | 25.0 | 95.3 | 67.2 | 56.3 | 84.4 | 38.9 | 41.8 | 30.3 | 71.4 | 65.4 | 20.0 |
| ES | 0.0 | 0.0 | 0.0 | 0.0 | 78.1 | 0.0 | 25.0 | 72.9 | 66.7 | 40.0 | 9.8 | 81.8 | 24.2 | 10.5 |
| Esb | 0.0 | 0.0 | 0.0 | 0.0 | 87.5 | 1.6 | 34.4 | 52.1 | 38.9 | 23.6 | 0.0 | 0.0 | 0.0 | 0.0 |
| GD | 0.0 | 5.0 | 0.0 | 25.0 | 78.1 | 15.6 | 50.0 | 49.0 | 63.9 | 34.5 | 0.5 | 100.0 | 0.8 | 0.3 |
| Gha | 80.0 | 98.8 | 71.7 | 50.0 | 95.3 | 81.3 | 93.8 | 95.8 | 83.3 | 69.1 | 22.5 | 55.2 | 85.3 | 36.8 |
| Gho | 16.7 | 62.5 | 11.7 | 25.0 | 84.4 | 26.6 | 43.8 | 45.8 | 61.1 | 34.5 | 3.5 | 66.7 | 24.2 | 3.8 |
| GL | 93.3 | 100.0 | 30.0 | 37.5 | 96.9 | 65.6 | 65.6 | 75.0 | 91.7 | 58.2 | 6.0 | 48.6 | 20.6 | 22.2 |
| GP | 40.0 | 95.0 | 55.0 | 37.5 | 98.4 | 62.5 | 78.1 | 89.6 | 97.2 | 78.2 | 26.6 | 52.1 | 60.6 | 29.8 |
| HN | 36.7 | 83.8 | 61.7 | 50.0 | 96.9 | 64.1 | 37.5 | 85.4 | 75.0 | 25.5 | 16.5 | 79.4 | 62.8 | 20.0 |
| JA | 70.0 | 85.0 | 61.7 | 37.5 | 100.0 | 82.8 | 78.1 | 63.5 | 80.6 | 61.8 | 30.1 | 64.7 | 41.8 | 10.8 |
| Jbo | 23.3 | 43.8 | 13.3 | 75.0 | 100.0 | 32.8 | 34.4 | 65.6 | 77.8 | 60.0 | 2.4 | 90.9 | 7.1 | 3.5 |
| JBr | 90.0 | 97.5 | 78.3 | 75.0 | 100.0 | 78.1 | 96.9 | 93.8 | 97.2 | 67.3 | 46.5 | 70.1 | 87.5 | 27.6 |
| JJ | 36.7 | 82.5 | 63.3 | 62.5 | 98.4 | 85.9 | 56.3 | 92.7 | 41.7 | 43.6 | 27.3 | 74.1 | 56.9 | 17.1 |
| JM | 83.3 | 100.0 | 63.3 | 50.0 | 100.0 | 82.8 | 100.0 | 82.3 | 94.4 | 76.4 | 51.5 | 67.5 | 58.3 | 25.4 |
| JM | 0.0 | 1.3 | 0.0 | 25.0 | 78.1 | 0.0 | 46.9 | 75.0 | 91.7 | 91.7 | 0.0 | 0.0 | 0.0 | 0.0 |
| JMf | 93.3 | 96.3 | 73.3 | 62.5 | 100.0 | 95.3 | 71.9 | 91.7 | 83.3 | 50.9 | 32.9 | 60.0 | 50.3 | 20.6 |
| JS | 50.0 | 82.5 | 41.7 | 50.0 | 93.8 | 68.8 | 81.3 | 96.9 | 100.0 | 65.5 | 50.1 | 47.0 | 100.0 | 100.0 |
| Jsa | 30.0 | 75.0 | 45.0 | 50.0 | 92.2 | 65.6 | 59.4 | 81.3 | 83.3 | 67.3 | 20.1 | 64.7 | 34.0 | 10.8 |
| JSb | 63.3 | 86.3 | 60.0 | 62.5 | 93.8 | 75.0 | 84.4 | 75.0 | 86.1 | 60.0 | 26.8 | 80.0 | 32.7 | 11.1 |
| JSc | 33.3 | 90.0 | 51.7 | 62.5 | 98.4 | 71.9 | 75.0 | 76.0 | 77.8 | 43.6 | 15.6 | 78.6 | 42.0 | 8.9 |
| JW | 33.3 | 65.0 | 40.0 | 87.5 | 96.9 | 64.1 | 90.6 | 85.4 | 88.9 | 61.8 | 8.1 | 83.3 | 40.8 | 5.7 |
| KA | 83.3 | 96.3 | 55.0 | 87.5 | 100.0 | 76.6 | 78.1 | 79.2 | 77.8 | 65.5 | 43.7 | 73.0 | 66.2 | 23.5 |
| KK | 33.3 | 56.3 | 8.3 | 0.0 | 93.8 | 40.6 | 46.9 | 81.3 | 100.0 | 76.4 | 9.6 | 54.1 | 34.4 | 19.4 |
| KL | 0.0 | 6.3 | 1.7 | 0.0 | 92.2 | 4.7 | 28.1 | 68.8 | 88.9 | 61.8 | 7.7 | 38.9 | 14.6 | 5.7 |
| KM | 0.0 | 0.0 | 0.0 | 0.0 | 53.1 | 0.0 | 43.8 | 69.8 | 66.7 | 46.6 | 0.0 | 0.0 | 0.0 | 0.0 |
| KS | 73.3 | 93.8 | 11.7 | 100.0 | 71.9 | 26.6 | 84.4 | 84.4 | 86.1 | 52.7 | 45.9 | 75.3 | 62.3 | 25.7 |
| KW | 0.0 | 3.8 | 0.0 | 50.0 | 95.3 | 0.0 | 84.4 | 82.3 | 80.6 | 50.9 | 1.3 | 75.0 | 8.5 | 1.3 |
| LH | 56.7 | 80.0 | 73.3 | 87.5 | 96.9 | 76.6 | 90.6 | 92.7 | 88.9 | 76.4 | 43.0 | 64.6 | 36.4 | 26.0 |
| LM | 16.7 | 27.5 | 1.7 | 0.0 | 67.2 | 4.7 | 28.1 | 57.3 | 61.1 | 32.7 | 9.4 | 40.0 | 12.7 | 3.2 |
| MAD | 60.0 | 95.0 | 75.0 | 62.5 | 96.9 | 81.3 | 81.3 | 88.5 | 83.3 | 63.6 | 17.6 | 87.0 | 23.8 | 7.3 |
| MD | 26.7 | 47.5 | 26.7 | 37.5 | 96.9 | 50.0 | 12.5 | 57.3 | 38.9 | 58.2 | 3.5 | 18.2 | 16.2 | 10.5 |
| MH | 0.0 | 13.8 | 5.0 | 25.0 | 100.0 | 6.3 | 59.4 | 69.8 | 80.6 | 74.5 | 51.9 | 65.5 | 47.3 | 17.5 |
| NC | 76.7 | 91.3 | 70.0 | 50.0 | 93.8 | 92.2 | 62.5 | 76.0 | 88.9 | 72.7 | 42.4 | 74.2 | 80.5 | 22.2 |
| NH | 86.7 | 100.0 | 85.0 | 62.5 | 98.4 | 93.8 | 90.6 | 86.5 | 94.4 | 70.9 | 32.8 | 90.9 | 52.6 | 7.0 |
| PB | 96.7 | 98.8 | 65.0 | 62.5 | 100.0 | 95.3 | 62.5 | 81.3 | 80.6 | 69.1 | 34.9 | 45.9 | 34.1 | 23.5 |
| PBL | 0.0 | 38.8 | 15.0 | 37.5 | 100.0 | 34.4 | 71.9 | 89.6 | 97.2 | 80.0 | 13.0 | 55.3 | 16.1 | 12.1 |
| PE | 13.3 | 45.0 | 10.0 | 25.0 | 96.9 | 20.3 | 50.0 | 79.2 | 80.6 | 41.8 | 32.0 | 52.7 | 64.4 | 64.4 |
| PM | 10.0 | 66.3 | 48.3 | 25.0 | 92.2 | 65.6 | 62.5 | 69.8 | 47.2 | 30.9 | 9.3 | 63.2 | 16.7 | 6.0 |
| PR | 56.7 | 85.0 | 46.7 | 75.0 | 100.0 | 60.9 | 87.5 | 83.3 | 80.6 | 50.9 | 9.3 | 64.0 | 24.0 | 7.9 |
| PW | 0.0 | 0.0 | 3.3 | 25.0 | 100.0 | 3.1 | 56.3 | 74.0 | 50.0 | 34.5 | 7.2 | 39.1 | 26.6 | 14.6 |
| Pwa | 33.3 | 70.0 | 35.0 | 50.0 | 95.3 | 65.6 | 68.8 | 87.5 | 80.6 | 45.5 | 6.9 | 78.9 | 24.2 | 6.0 |
| RH | 6.7 | 21.3 | 1.7 | 25.0 | 82.8 | 3.1 | 62.5 | 89.6 | 83.3 | 61.8 | 44.6 | 50.7 | 96.8 | 64.4 |
| RL | 13.3 | 61.3 | 53.3 | 62.5 | 96.9 | 85.9 | 62.5 | 92.7 | 80.6 | 72.7 | 28.3 | 77.5 | 42.5 | 12.7 |
| RR | 10.0 | 51.3 | 18.3 | 25.0 | 98.4 | 42.2 | 56.3 | 82.3 | 88.9 | 43.6 | 13.9 | 67.7 | 34.8 | 9.8 |
| SH | 6.7 | 48.8 | 23.3 | 25.0 | 84.4 | 51.6 | 43.8 | 69.8 | 55.6 | 49.5 | 22.0 | 53.4 | 59.7 | 23.2 |
| SL | 0.0 | 0.0 | 0.0 | 0.0 | 50.0 | 0.0 | 34.4 | 52.1 | 88.9 | 52.7 | 0.0 | 0.0 | 0.0 | 0.0 |
| ST | 50.0 | 80.0 | 50.0 | 25.0 | 92.2 | 79.7 | 71.9 | 88.5 | 50.0 | 38.2 | 9.4 | 66.7 | 33.6 | 9.5 |
| TA | 10.0 | 35.0 | 11.7 | 12.5 | 75.0 | 35.9 | 34.4 | 42.7 | 33.3 | 52.7 | 6.6 | 64.3 | 23.8 | 4.4 |
| TJ | 93.3 | 98.8 | 88.3 | 62.5 | 98.4 | 92.2 | 68.8 | 88.5 | 50.0 | 52.7 | 24.9 | 66.7 | 56.0 | 16.2 |
| TK | 0.0 | 35.0 | 6.7 | 37.5 | 100.0 | 0.0 | 87.5 | 71.9 | 69.4 | 54.5 | 20.8 | 60.6 | 48.4 | 21.0 |
| WC | 30.0 | 0.0 | 45.0 | 62.5 | 96.9 | 57.8 | 75.0 | 94.8 | 52.8 | 70.9 | 46.1 | 56.6 | 86.6 | 38.7 |
| WE | 46.7 | 73.8 | 51.7 | 62.5 | 100.0 | 79.7 | 84.4 | 87.5 | 91.7 | 70.9 | 26.1 | 59.4 | 58.9 | 21.9 |
| WM | 36.7 | 55.0 | 25.0 | 37.5 | 92.2 | 35.9 | 50.0 | 61.5 | 61.1 | 43.6 | 3.5 | 57.1 | 15.3 | 4.4 |

Supplementary Table S6: Individual deficits of auditory processing in participants with post-stroke aphasia

Matrix showing results of Bayesian Test for a Deficit, controlling for years of education as a covariate, comparing each participant with aphasia’s (rows) performance on each of seven psychoacoustic tests (columns) to the control group. Each cell contains the uncorrected, one-sided p-value for a significance test of whether the participant with aphasia (indicated in the left-most cell of that row) has a score on the psychoacoustic test (indicated in the top-most cell of that column) that is an observation from the control group, controlling for years of education as a covariate. Participants are ordered alphabetically. Abbreviations: NA = Not Acquired as patient unable to perform this psychoacoustic task at the easiest difficulty level. * indicates the p-value is significant at the Bonferroni corrected significance threshold of p<0.007 (corrected for 7 comparisons) (significant cells are also shaded in green).

| **Participant with aphasia** | **Log_10_ pitch basic change detection threshold** | **Pitch detection of local change** | **(Pitch detection of global change)^4^** | **Log_10_ rhythm single time interval discrimination threshold** | $\boldsymbol{\surd}$**(Rhythm isochrony deviation detection threshold)** | **Rhythm metrical pattern discrimination threshold** | **Log_10_ Dynamic Modulation detection threshold** |
| --- | --- | --- | --- | --- | --- | --- | --- |
| **AG** | 0.08 | 0.04 | 0.15 | 0.11 | 0.19 | 0.12 | 0.17 |
| **AL** | 0.21 | 0.08 | 0.42 | 0.20 | 0.03 | 0.69 | 0.07 |
| **BH** | 0.11 | 0.06 | 0.02 | 0.05 | 0.17 | 0.17 | **0.006*** |
| **CH** | 0.10 | 0.81 | 0.53 | 0.64 | 0.55 | 0.17 | 0.17 |
| **DF** | **0.002*** | 0.04 | 0.22 | **NA** | **NA** | **NA** | 0.01 |
| **DM** | 0.87 | 0.19 | 0.15 | 0.09 | 0.39 | 0.13 | 0.51 |
| **Ebo** | 0.61 | 0.64 | 0.29 | 0.07 | 0.27 | 0.23 | 0.19 |
| **GP** | 0.12 | 0.24 | 0.15 | 0.20 | 0.27 | 0.02 | **0.0003*** |
| **JS** | 0.26 | 0.43 | 0.74 | 0.86 | 0.20 | 0.70 | 0.28 |
| **MAD** | 0.21 | 0.01 | 0.02 | 0.01 | **0.0005*** | 0.17 | **0.002*** |
| **MH** | **0.007*** | 0.31 | 0.29 | 0.03 | 0.11 | 0.31 | **0.0006*** |
| **NC** | 0.10 | 0.01 | 0.11 | 0.72 | 0.46 | 0.82 | 0.45 |
| **PBL** | 0.25 | 0.09 | 0.07 | 0.05 | **NA** | 0.06 | 0.12 |
| **PR** | **0.001*** | 0.02 | 0.02 | **0.004*** | **NA** | **NA** | **0.0004*** |
| **RH** | 0.56 | 0.80 | 0.73 | 0.69 | 0.78 | 0.66 | 0.04 |
| **ST** | 0.02 | 0.02 | 0.01 | 0.07 | **0.006*** | 0.31 | 0.01 |
| **WE** | 0.01 | 0.22 | 0.22 | 0.05 | 0.06 | 0.41 | 0.11 |

Supplementary Table S7: Correlations between Principal Component 3 and neuropsychological tests

‘P value’ corresponds to uncorrected one-sided p-values from Spearman correlations between neuropsychological scores and Principal Component 3 score in the post-stroke aphasia subgroup who underwent psychoacoustic testing. * indicates the p-value is significant at the Bonferroni corrected significance threshold of p<0.004 (corrected for 14 comparisons). Abbreviation: ‘CSB’ = Cambridge Semantic Battery.

| **Neuropsychological test** | **Spearman’s rho with Principal Component 3** | **P value** |
| --- | --- | --- |
| ‘Cookie Theft’ number of speech tokens | 0.92 | 0.00000006* |
| ‘Cookie Theft’ Mean Length of Utterance | 0.91 | 0.0000003* |
| ‘Cookie Theft’ Words Per Minute | 0.91 | 0.0000002* |
| Immediate non-word repetition | -0.11 | 0.34 |
| Immediate word repetition | 0.01 | 0.49 |
| Boston Naming Test | -0.24 | 0.18 |
| CSB Picture Naming | -0.06 | 0.41 |
| Forward Digit Span | -0.30 | 0.12 |
| CSB Spoken Word-to-Picture Matching | -0.33 | 0.10 |
| Spoken Sentence Comprehension | -0.30 | 0.13 |
| Synonym Judgement Test | 0.18 | 0.25 |
| ‘Cookie Theft’ Type-Token Ratio | -0.27 | 0.15 |
| Raven’s Coloured Progressive Matrices | 0.34 | 0.09 |
| Brixton Spatial Anticipation Test | 0.19 | 0.23 |

Additional methodological details

Participants

The PSA subgroup were recruited from the larger cohort of 76 stroke survivors on the basis of willingness and availability to undergo further psychoacoustic testing.

Lesion overlap map

Structural T_1_-weighted MRI scans were acquired on a 3.0 T Philips Achieva scanner (Philips Healthcare) using an 8-element SENSE head coil. A T_1_-weighted inversion recovery sequence with 3D acquisition was used with the following parameters: repetition time = 9.0ms, echo time = 3.93ms, flip angle = 8°, 150 contiguous slices, slice thickness = 1mm, acquired voxel size = 1.0 x 1.0 x 1.0mm, matrix size 256 x 256, field of view = 256mm, inversion time = 1150ms, SENSE acceleration factor 2.5, total scan acquisition time = 575s. The lesion overlap map was generated using FSL’s ‘fslmaths’ command (Jenkinson *et al.*, 2012) and displayed in MRIcron (Rorden and Brett, 2000) (www.mricro.com/mricron/).

Neuropsychological tests

The entire cohort of 76 stroke survivors had previously been administered an extensive battery of neuropsychological tests and principal component analysis (PCA) results from this database have been published from earlier data collection rounds (Butler *et al.*, 2014; Halai *et al.*, 2017). We expected immediate non-word and word repetition (subtests 8 and 9 from the Psycholinguistic Assessment of Language Processing in Aphasia battery (Kay *et al.*, 1992)), Boston Naming Test (BNT) (Kaplan *et al.*, 1983), Forward Digit Span (Wechsler, 1987), and Cambridge Semantic Battery picture naming (Bozeat *et al.*, 2000) to represent participants’ phonological ability. We expected the Cambridge Semantic Battery spoken word-to-picture matching (Bozeat *et al.*, 2000), spoken sentence comprehension from the Comprehension Aphasia Test (CAT) (Swinburn *et al.*, 2005), 96-trial synonym judgement test (Jefferies *et al.*, 2009) and Type-Token Ratio from the ‘Cookie theft’ description (Boston Diagnostic Aphasia Examination (Goodglass and Kaplan, 1983)) to represent participants’ semantic ability. We expected the number of speech tokens, Mean Length of Utterance, and Words Per Minute from the ‘Cookie theft’ description (Boston Diagnostic Aphasia Examination (Goodglass and Kaplan, 1983)) to represent participants’ speech fluency. We expected the Raven’s Coloured Progressive Matrices (Raven, 1962) and the Brixton Spatial Anticipation Test (Burgess and Shallice, 1997) to represent participants’ executive function.

The ‘Cookie theft’ description from the Boston Diagnostic Aphasia Examination (Goodglass and Kaplan, 1983) involved being shown the ‘Cookie theft’ picture and being asked ‘tell me everything you see going on in this picture’. Responses were recorded, transcribed and used to obtain various parameters of connected speech production (Type-Token Ratio, number of speech tokens, Mean Length of Utterance, Words Per Minute) using the coding method described previously (Halai *et al.*, 2017).

Psychoacoustic tests of pitch, rhythm and timbre

Psychoacoustic testing occurred over several sessions for each participant with PSA, with the number of sessions determined by the participant so as to avoid fatigue impacting on performance. With the controls, psychoacoustic testing was always completed within a single session. Due to time constraints, one control was not tested on P2 or P3, and a second healthy control was not tested on P1.

We define tone sequences as more than two tones grouped together. Thus, four of the seven psychoacoustic tests required processing of tone sequences: P2 (pitch local change detection); P3 (pitch global change detection); R2 (isochrony deviation detection); and R3 (metrical pattern discrimination). Three of the seven psychoacoustic tests did not require processing of tone sequences: P1 (pitch basic change detection); R1 (single time interval discrimination); and DM (Dynamic Modulation detection).

Statistical analysis

All variables were assessed for normality using the Kolmogorov-Smirnov test.

Group differences between PSA and control participants were initially assessed using the independent samples *t*-test for age (normally distributed), Pearson’s Chi-square test for sex, and Mann-Whitney U test for years of education (not normally distributed; p=0.00002 in PSA). As several pure tone audiometry thresholds were not normally distributed, group differences of audiometry thresholds between PSA and control participants were assessed using Mann-Whitney U tests.

As the PSA subgroup and control group had significantly different years of education (see Results), and years of education was not normally distributed, group differences between PSA and control participants on neuropsychological and psychoacoustic measures were assessed using a non-parametric one-way rank analysis of covariance (ANCOVA) with years of education included as the covariate (Quade, 1967).

It was possible that participants with PSA might have been significantly impaired on psychoacoustic tasks at the individual level, even if there was no group difference compared to controls. We therefore compared, individually, the psychoacoustic scores of each participant with PSA to the control group using the Bayesian Test for a Deficit controlling for years of education as a covariate (https://homepages.abdn.ac.uk/j.crawford/pages/dept/SingleCaseMethodsComputerPrograms.HTM) (Crawford *et al.*, 2011). This method provides a Bayesian point estimate of the proportion of controls expected to perform worse than the individual case, which corresponds to the one-sided p-value for a significance test of whether the case’s score can be assumed to equal an observation from the control population (Crawford and Garthwaite, 2006). Since this method assumes a Gaussian distribution, psychoacoustic variables that were not normally distributed in either the PSA or control group (P1, P3, R1, R2, DM) were log_10_ transformed. If the log_10_ transformed variable was not normally distributed, alternative transformations were used. R2 only became normally distributed in controls after square root transformation. P3 only became normally distributed in controls after quartic transformation. P1 remained non-normally distributed in the control group, no matter the transformation applied (p=0.01 for log_10_P1).

Given the large number of neuropsychological measures available for our PSA subgroup and the whole cohort we recruited them from, we used a varimax-rotated PCA to reduce these scores to a smaller number of dimensions, as has been done previously (Butler *et al.*, 2014; Halai *et al.*, 2017). As PCA was unlikely to be stable in our subgroup of 17 PSA participants with psychoacoustic data (Preacher and MacCallum, 2002), we performed the PCA on the correlation matrix of neuropsychological test scores of the entire cohort of PSA participants (n=76), which has been shown formally to be highly reliable and stable. Scores from Principal Components (PCs) with an eigenvalue greater than 1 were taken to be estimates of underlying cognitive components in our 17 PSA participants and were used in correlation analyses with the psychoacoustic measures.

Within the PSA subgroup, correlations were performed between neuropsychological test scores, neuropsychological ‘PC’ scores and psychoacoustic scores. Non-parametric Spearman correlations were performed because several variables (P1, R2, DM, PC1) were not normally distributed.

We used SPSS version 25 for the above statistical analyses, except for partial Spearman correlations and rank ANCOVAs, which were performed in Matlab R2018a. We defined statistical significance as p<0.05 with Bonferroni correction (by the number of tests) applied to the significance thresholds (i.e. reported p-values are uncorrected). Reported p-values for correlations between neuropsychological and/or psychoacoustic scores are one-tailed due to *a priori* hypotheses as to the direction of the associations, i.e. that better performance on auditory tasks would be associated with higher language scores.
